# The melissopalynological investigation in the Eastern Dry Zone of Karnataka, India

**DOI:** 10.1101/2020.07.06.189274

**Authors:** D. Shishira, A. R. Uthappa, Veeresh Kumar, Shringeshwara, G. C. Kuberappa

## Abstract

Melissopalynology, the analysis of pollen grains present in honey, indicates about the pollen and nectar sources in a region utilized by bees, which is used to determine the bee floral resources and botanical origin of the honey. This study investigated the melissopalynological analysis of the honey samples from the Eastern Dry zone of Karnataka. 24 honey samples were examined based on pollen analyses, among them 14 samples were unifloral, rest were multifloral. The unifloral honey had pollens of *Callistemon viminalis, Areca catechu, Citrus* sp., *Mallotus philippensis, Cocos nucifera, Eucalyptus* sp., *Ocimum* sp., *Moringa oleifera* and *Pongamia pinnata*. Samples collected in October, November, December, and January were rich in pollens of *Eucalyptus* sp.. Similarly, samples collected in January, February and March had pollen of tree species *viz*., *Swietenia mahagoni, Canthium parviflorum, Simarouba glauca, Eucalyptus* sp., *Moringa oleifera, Syzygium cumini, Tabebuia* sp., *Pongamia pinnata, Acanthaceae, Anacardium occidentale, Cocos nucifera, Areca catechu, Mallotus philippensis, Bauhinia variegata, Psidium guajava, Alangiaceae, Euphorbiaceae, Ulmaceae, Capparis zeylanica, Convolvulaceae*. GKVK-11 followed by GKVK-12 sample recorded the highest Shannon diversity and GKVK-9 followed by GKVK-7 sample recorded the least diversity. Based on the similar floral composition samples were classified into four clusters. The PCA revealed that most of the samples grouped into a single cluster, except 7, 19, 20, 21, and 22 which were placed away from the origin. The presence of pollen in the honey of a particular plant species during different months is related to the blooming of that particular plant species from which the bees forage. The flora of honey changes with the season. The diversity of pollen grains in honey varied with location to location. The present study provides scientific knowledge to the beekeepers by indicating important plants for the development of the regional apiculture, through the identification of pollen types.

## Introduction

Melissopalynology is a study of pollen grains present in honey. It helps to assess the bee floral availability of the region, where the particular types of honey are produced (Sajwani *et al*., 2007; Song *et al*., 2010). It also provides the exact information regarding the floral resource to bees in the vicinity and also becomes useful in the construction of the floral calendar (Sekhar, 2000).

The honey bees depend on the flora for nectar and pollen. The quality of honey varies concerning the flora available in and around them (Ponnuchamy *et al*., 2014). The Eastern dry zone of Karnataka is one of the important areas for beekeeping since the region comprises of a rich diversity of bee flora. One or the other flora bloom in every month which makes the bees survive throughout the year. The relative abundance of bee plants in the eastern dry zone consisted of trees (46%), herbs (25%), shrubs (18%), trailers (9%), climbers, and palms (2%) (Marc, 2012). The major bee flora available in and around the eastern dry zone region was *Callistemon viminalis* Byrnes, *Anacardium occidentale* L., *Antigonon leptopus* Hook. & Arn., *Simarouba glauca* DC., *Eucalyptus* sp., *Tabebuia* sp. L., *Cocos nucifera* L., *Mallotus philippensis* Lour, *Santalum album* L., *Tamarindus indica* L., *Trewia nudiflora* L., and *Bauhinia purpurea* L. (Sekhar, 2000).

The urban beekeeping industry is gaining popularity and is accepted as a complementary activity to agriculture in India (Attri, 2010). The analysis of the honey pollen spectrum is extremely useful to detect the contribution of different nectar sources during a different period of the year (Oliviera *et al*., 2010). It contributes to the development of the beekeeping industry by the way of proper utilization of floral resources and provides the exact information about the floral resources of the bees in the particular region. Melissopalynological studies are least touched in various parts of India, especially in South India (Jhansi *et al*., 1994). The studies on the source of pollen have been very few in Asia and with respect to different species of honey bees, the melissopalynological information available is very less (Ramalho *et al*., 2007 and Novais *et al*., 2009).

Although honey has been the most widespread bee product, the bee pollen trade is undergoing expressive growth. The quantity and quality of the bee pollen produced have attracted increased attention for Indian Apiculture. Eastern dry zone of Karnataka needs in-depth studies on the bee floras available also to support the interest of the urban beekeepers related to when is the honey flow season, dearth period, when to feed bees, when not to feed bees for managing the colonies. It also helps in developing the pollen spectrum through pollen combination. Henceforth, to encourage them and to make them aware of the floral calendar and the flora to which bee visits. This study aimed to identify the botanical origin of pollen loads and the melissopalynological investigation of *Apis cerana* honey in the Eastern dry zone of Karnataka.

## Materials and Methods

### Honey sampling and Melissopalynology

Twenty-one honey samples (Tab.1) of *Apis cerana* were collected from different regions of the Eastern dry zone (Fig. 1) during 2016-17. A piece of *Apis cerana* comb completely sealed with honey was collected from different places and brought to the laboratory for extraction. The cut comb was unsealed using the uncapping knife, honey was extracted using a honey extractor, and the obtained honey thus filtered into a beaker using a muslin cloth and was transferred to an airtight container and was labeled concerned to the place, date and time of collection. The samples were stored in a refrigerator at 4°C for further studies (Method adopted from Saxena *et al*., 2010). After the complete removal of honey, the frame along with an empty comb was placed back to the same colony and the same position.

**Table 1.** List of samples examined and its location

| Sample No | Locality | Sample code | GPS Coordinates | Time of collection |
| --- | --- | --- | --- | --- |
| 1 | Baglur | Baglur-1 | 12.8304°N, 77.8662°E | July 2016 |
| 2 | Baglur | Baglur-2 | 12.8304°N, 77.8662°E | July 2016 |
| 3 | GKVK | GKVK-1 | 13.0797°N, 77.5808°E | July 2016 |
| 4 | GKVK | GKVK-2 | 13.0797°N, 77.5808°E | August 2016 |
| 5 | GKVK | GKVK-3 | 13.0797°N, 77.5808°E | August 2016 |
| 6 | Yelahanka | Yelahanka-1 | 13.1155°N, 77.6070°E | September 2016 |
| 7 | GKVK | GKVK-4 | 13.0797°N, 77.5808°E | September 2016 |
| 8 | GKVK | GKVK-5 | 13.0797°N, 77.5808°E | October 2016 |
| 9 | Baglur | Baglur-3 | 12.8304°N, 77.8662°E | October 2016 |
| 10 | M S Palya | M S Palya-1 | 13.0813°N, 77.5483°E | November 2016 |
| 11 | GKVK | GKVK-6 | 13.0797°N, 77.5808°E | November 2016 |
| 12 | GKVK | GKVK-7 | 13.0797°N, 77.5808°E | December 2016 |
| 13 | GKVK | GKVK-8 | 13.0797°N, 77.5808°E | December 2016 |
| 14 | GKVK | GKVK-9 | 13.0797°N, 77.5808°E | December 2016 |
| 15 | GKVK | GKVK-10 | 13.0797°N, 77.5808°E | December 2016 |
| 16 | Baglur | Baglur-4 | 12.8304°N, 77.8662°E | December 2016 |
| 17 | M S Palya | M S Palya-2 | 13.0813°N, 77.5483°E | January 2017 |
| 18 | Yelahanka | Yelahanka-2 | 13.1155°N, 77.6070°E | February 2017 |
| 19 | GKVK | GKVK-11 | 13.0797°N, 77.5808°E | March 2017 |
| 20 | Baglur | Baglur-5 | 12.8304°N, 77.8662°E | March 2017 |
| 21 | GKVK | GKVK-12 | 13.0797°N, 77.5808°E | March 2017 |
| 22 | GKVK | GKVK-13 | 13.0797°N, 77.5808°E | April 2017 |
| 23 | GKVK | GKVK-14 | 13.0797°N, 77.5808°E | May 2017 |
| 24 | GKVK | GKVK-15 | 13.0797°N, 77.5808°E | June 2017 |

**Figure 1.**
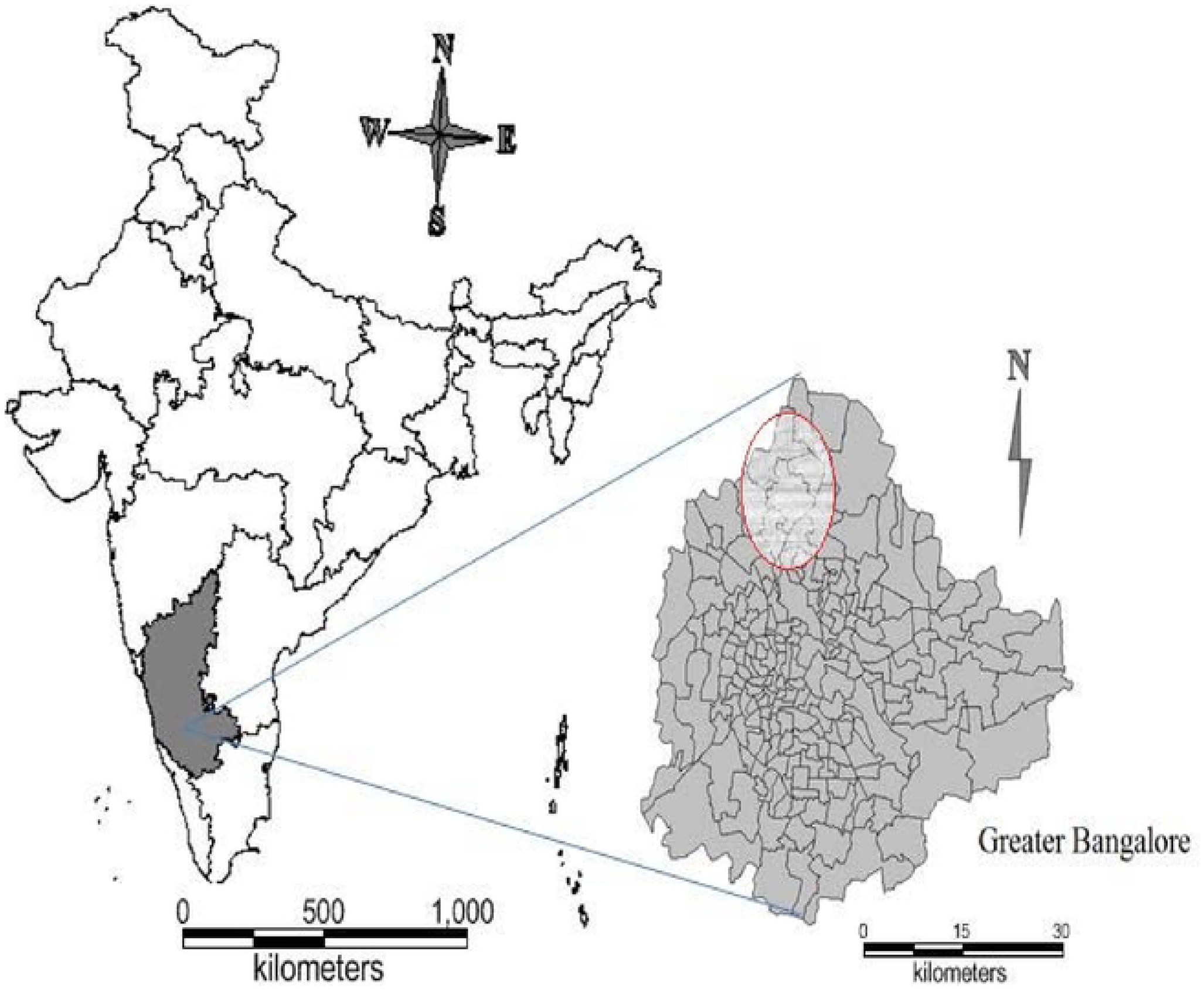
Location map showing the study area. (Left) map showing the position of Karnataka, (Right) map showing the sampling sites in the Bangalore region of Karnataka.

*Apis cerana* honey was analyzed for the presence of pollen using the acetolysis method suggested by Erdtman (1952, 1966). One gram of honey was diluted in 9ml of water and centrifuged at 4000rpm for 15 min. The pollen so obtained at the bottom was collected, identified, and counted in all the 25 cells of hemocytometer under a microscope (Motic). Based on the percentage of each pollen type present in the honey was classified as unifloral (more than 45% of single pollen) or multifloral (less than 45% of single pollen) (Chaturvedi, 1989).

The grouping of honey is done by taking into account of pollen grains in 10g honey following Maurizio’s classes. Group I (20,000 pollen grains per 10 g honey), Group II (20,000–100,000 grains per 10 g honey), Group III (100,000– 500,000 grains per 10 g honey) (Louveaux *et al*., 1978).

The images of pollen collected from honey samples were captured by Moticam 2300 3.0M Pixel USB2.0 and the software used was Motica image plus 2.0ML. These pollen images were identified by making slides of available flora.

To study the similarity between the pollen sources agglomerative hierarchical clustering method was run using XLSTAT. The dendrogram was constructed using the Jaccard co-efficient and Unweighted pair group average method. PAST 3.24 version was used for principal component analysis (PCA) of the pollen data.

## Results

Out of 24 samples analyzed, 15 were unifloral (62.5%) with predominate pollen viz., *Areca catechu* L., *Citrus* sp., *Mallotus philippensis* Lour., *Cocos nucifera* L., *Pongamia pinnata* L., *Ocimum* sp., *Casuarina equisetifolia* L. each count 4.17% and *Eucalyptus* sp. alone was 33.33%. Seven samples were identified as multifloral (37.50%) (Figure 3).

**Figure 2:**
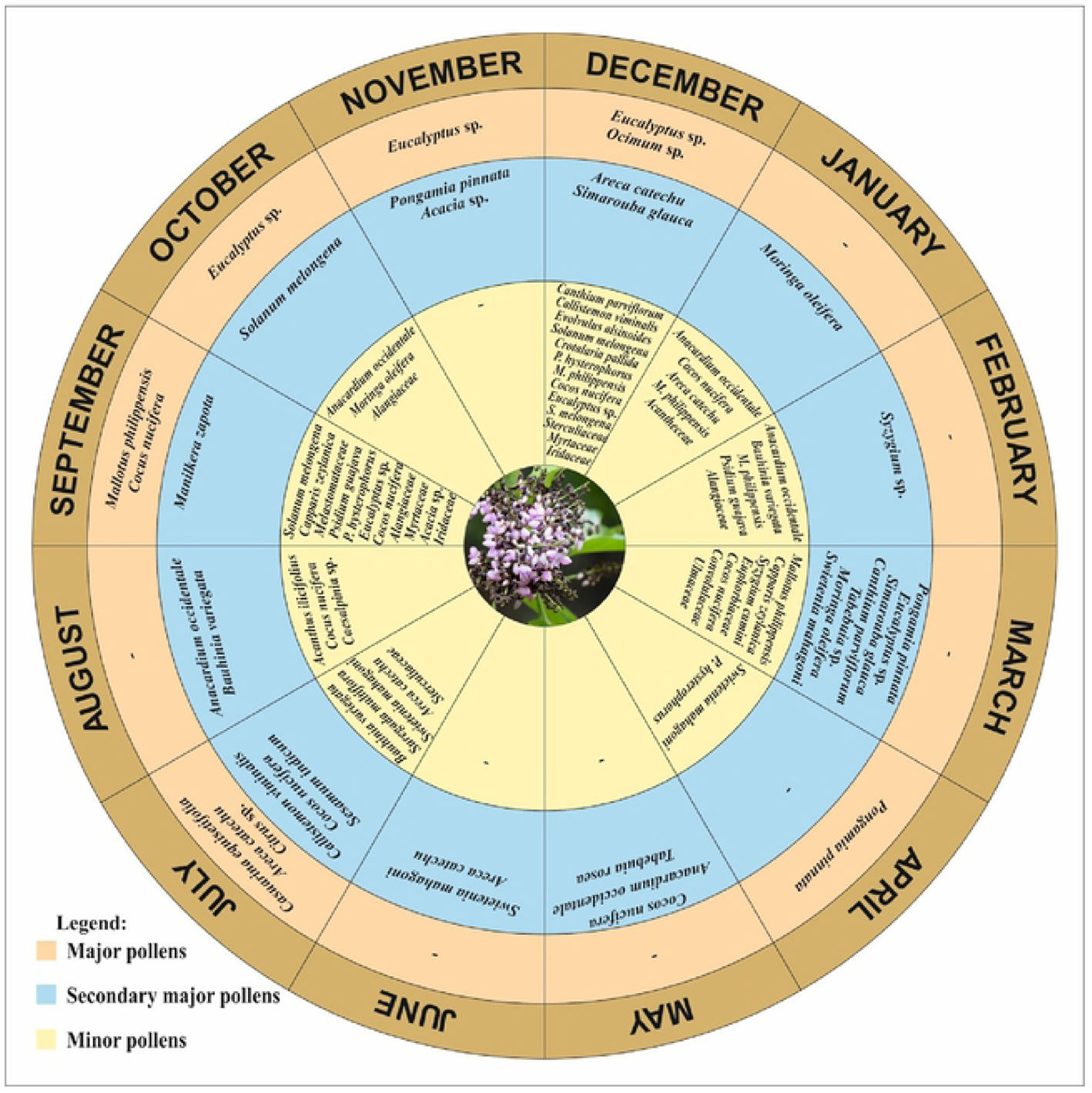
Floral Calender of *A*.*cerana* Fab. in Eastern Dry Zone of Karnataka.

**Figure 3:**
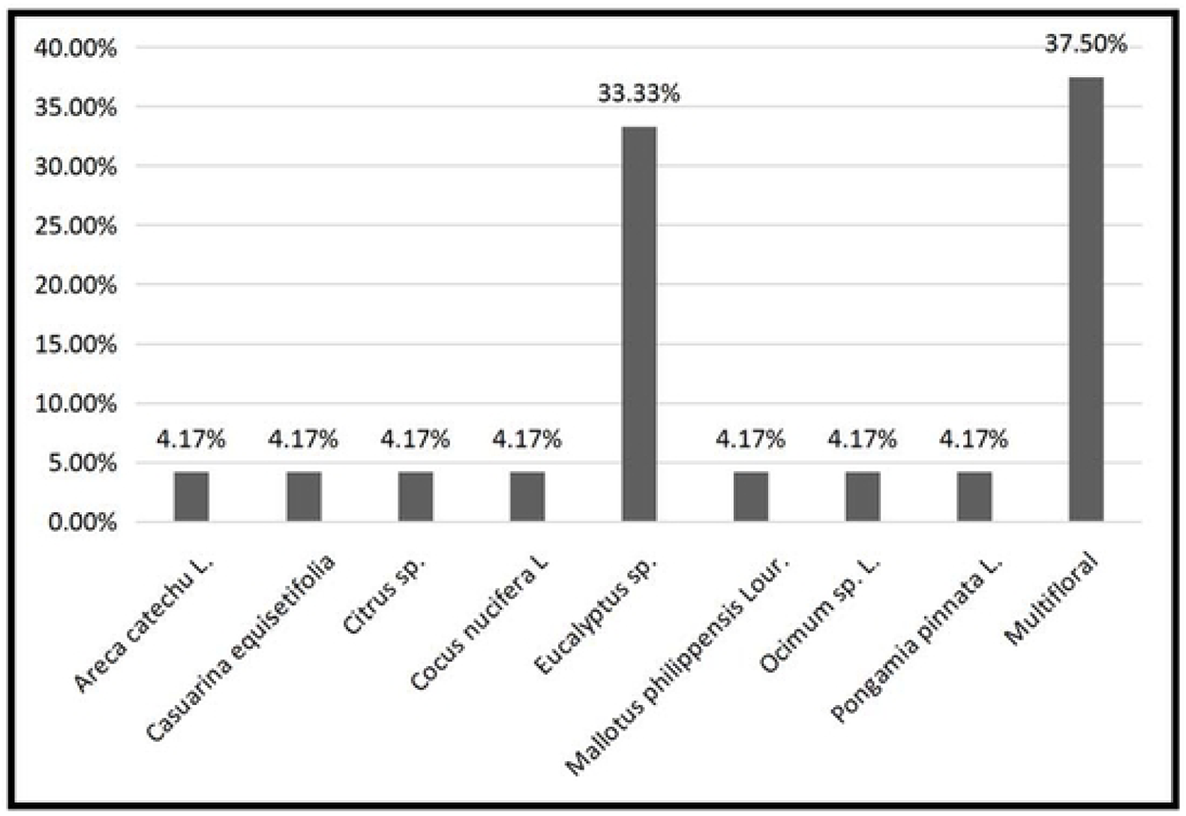
Floral nature of honey samples of *Apis cerana* collected from the Eastern *dry* zone of Karnataka are *Areca catechu, Citrus* sp., *Mallotus philippensis, Cocus nucifera, Simarouba glauca, Ocimum* sp., *Casuarina equisetifolia* each counts 4.17%, *Eucalyptus* sp. counts 33.33% and 9 samples were multifloral (37.50%)

The predominate families found in the samples are Acanthaceae, Convolvulaceae and Solanaceae(7.41% each), Alangiaceae, Anacardiaceae, Arecaceae, Asteraceae, Bignoniaceae, Capparaceae, Casuarinaceae, Fabaceae, Iridaceae, Lamiaceae, Melastomataceae, Meliaceae, Moringaceae, Pedaliaceae, Piperaceae, Rubiaceae, Rutaceae, Sapotaceae, Simaroubaceae, Sterculiaceae and Ulmaceae(3.70% each), Euphorbiaceae and Leguminosae (14.81% each) and Myrtaceae count to 18.52% (Figure 4). The absolute pollen counts per 10 g of honey samples indicated that 50% of honey samples belong to the group I (2,500-15,00 pollen grains), 37.50% of samples belong to group II (25,000-60,000) and 12.5% of samples belong to group III (1,20,000-1,42,5000) (Table 2, Figure 5).

**Table 2.**
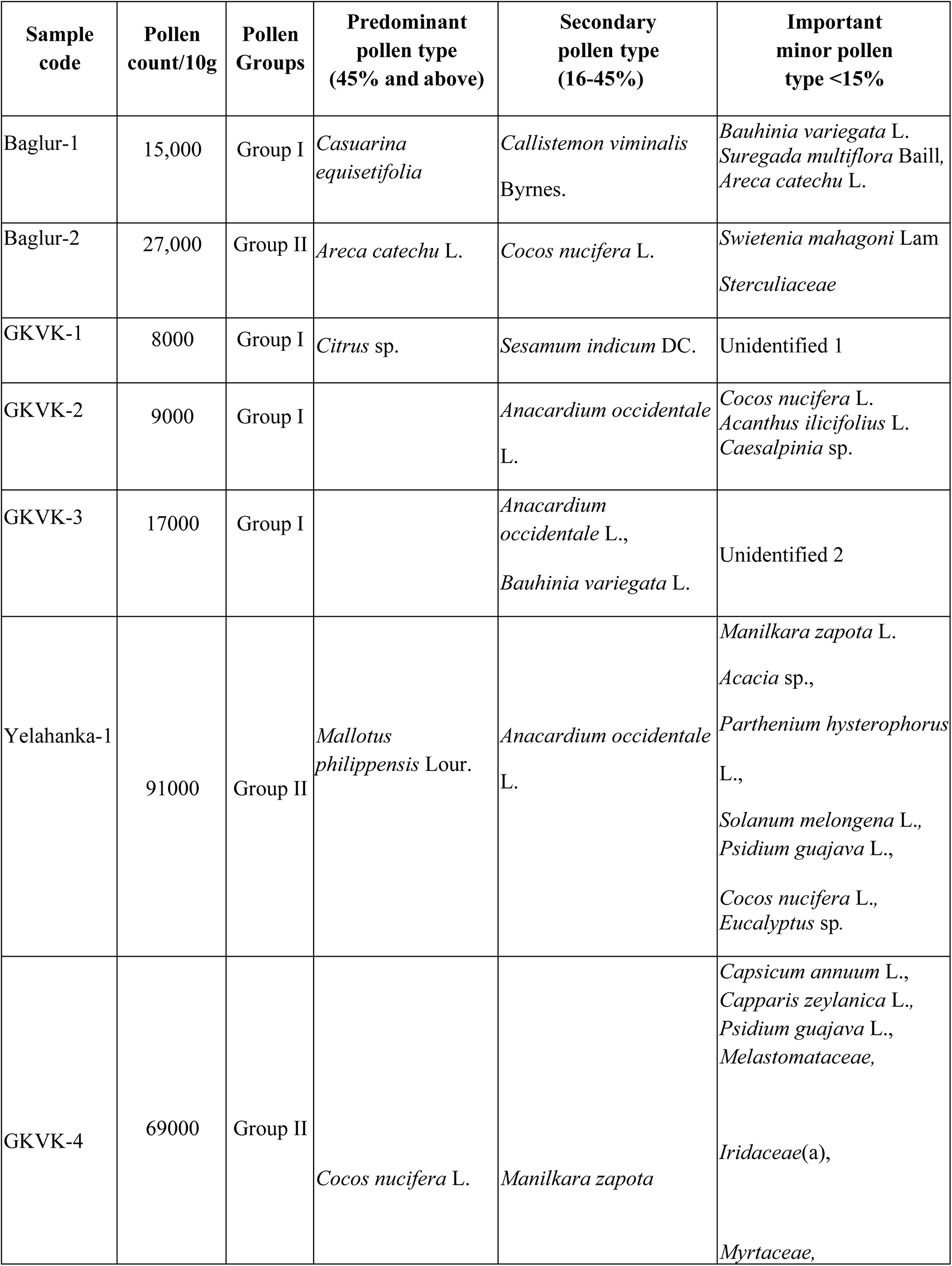
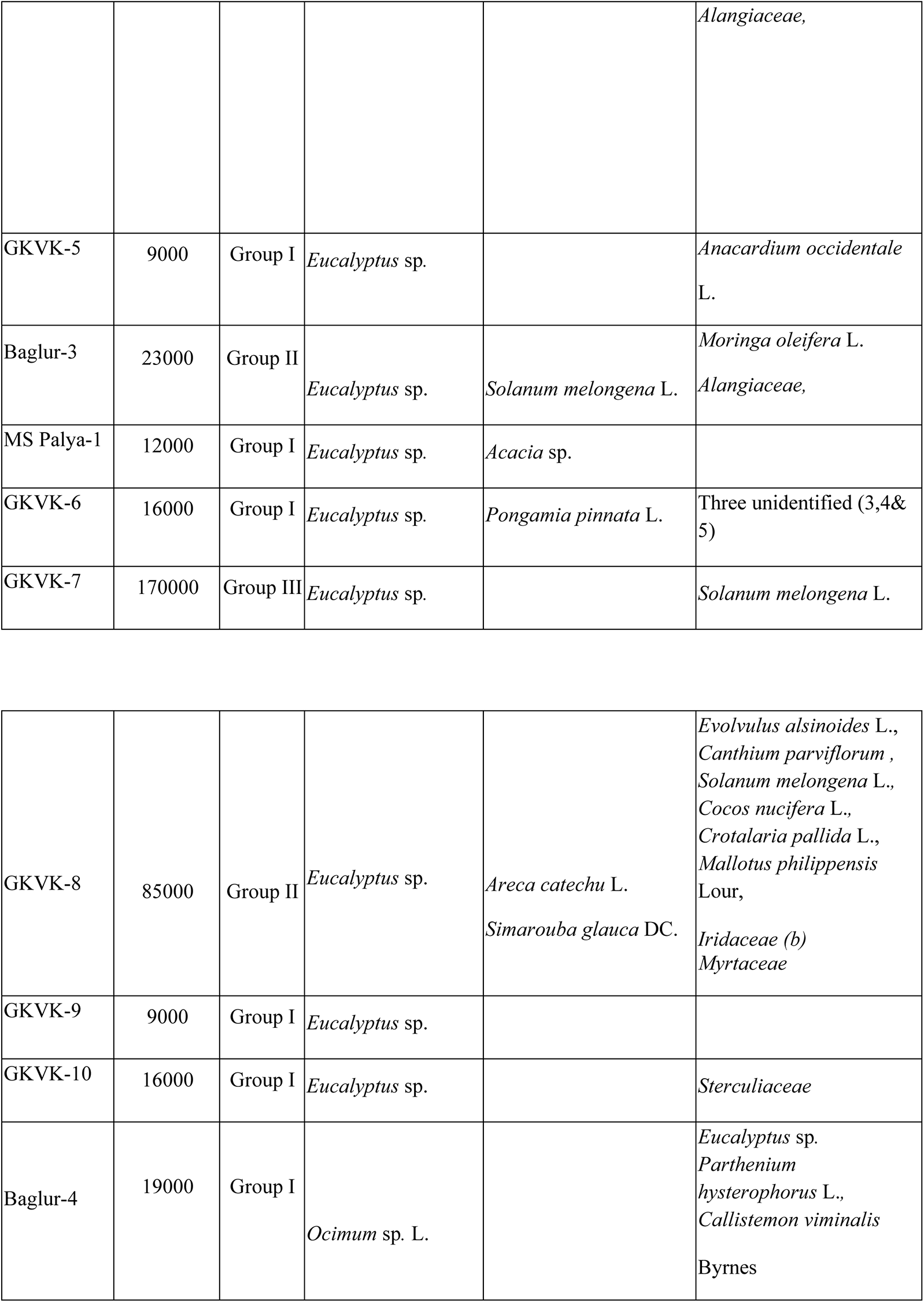
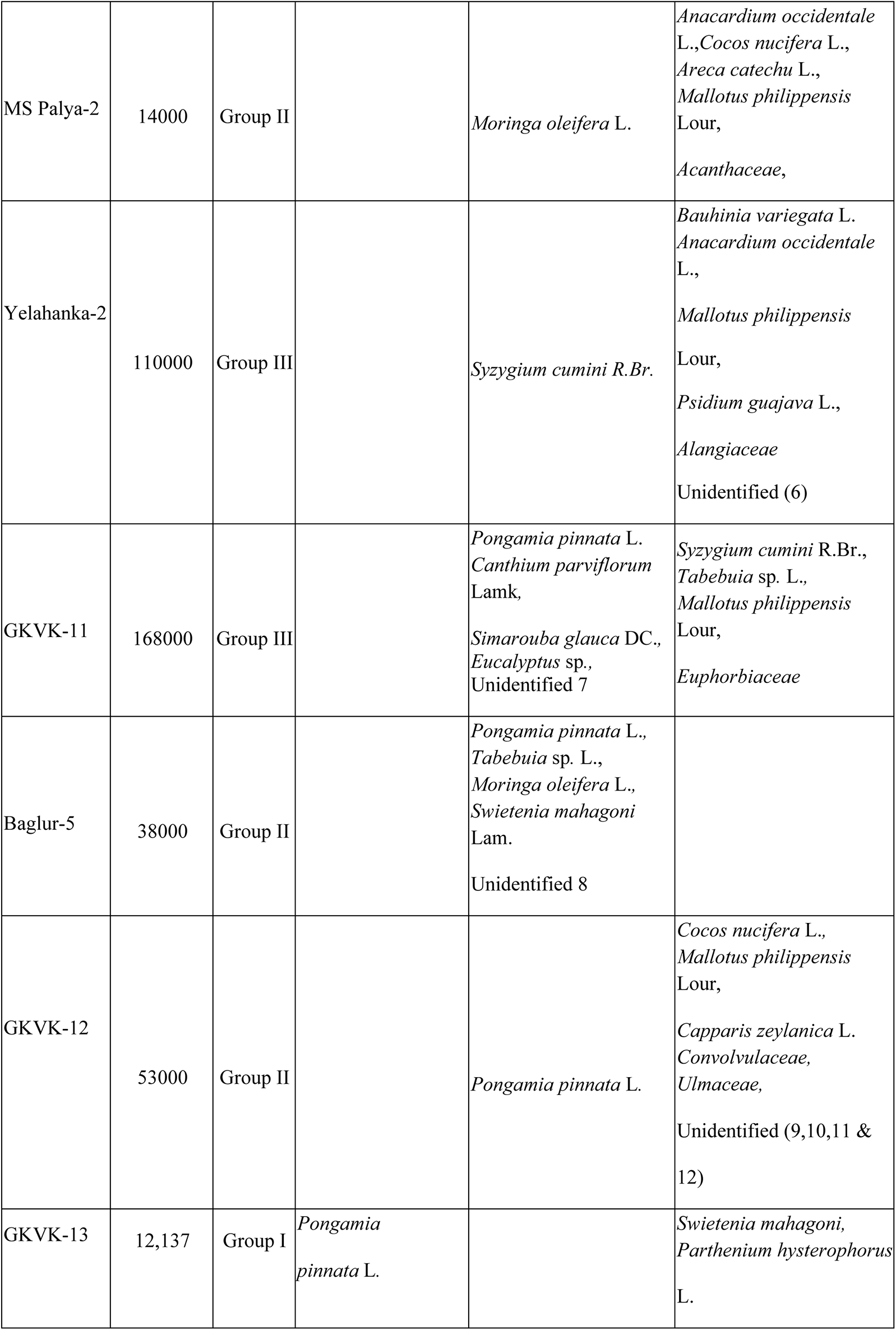
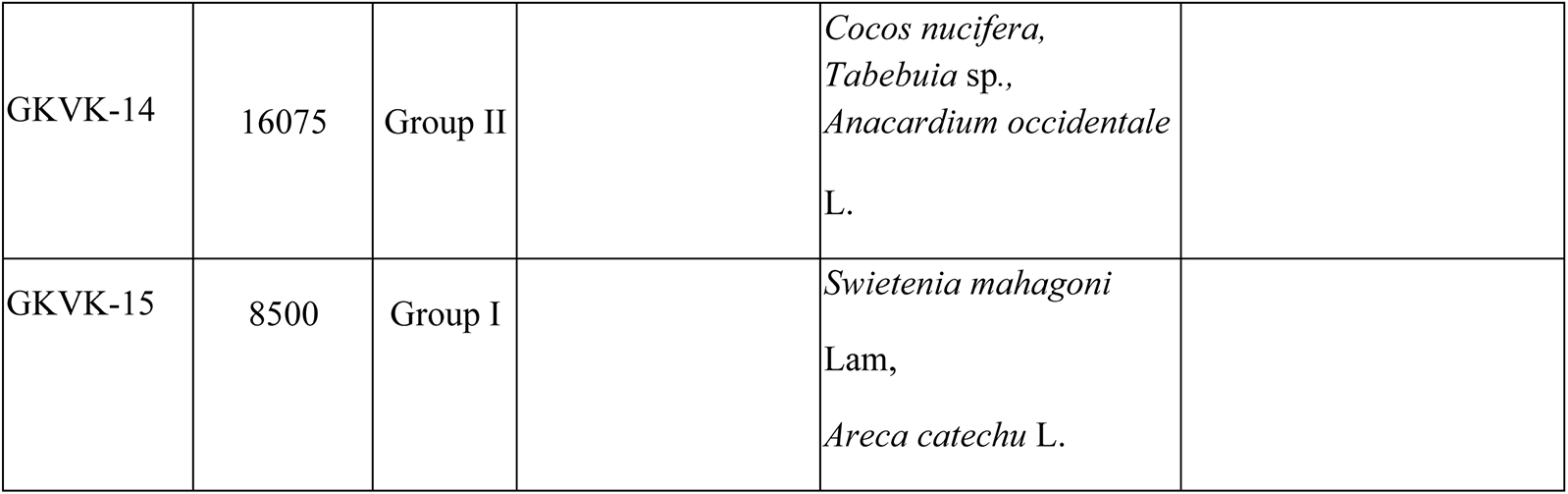
Melissopalynology data of honey samples collected in Eastern Dry region of Karnataka

**Figure 4:**
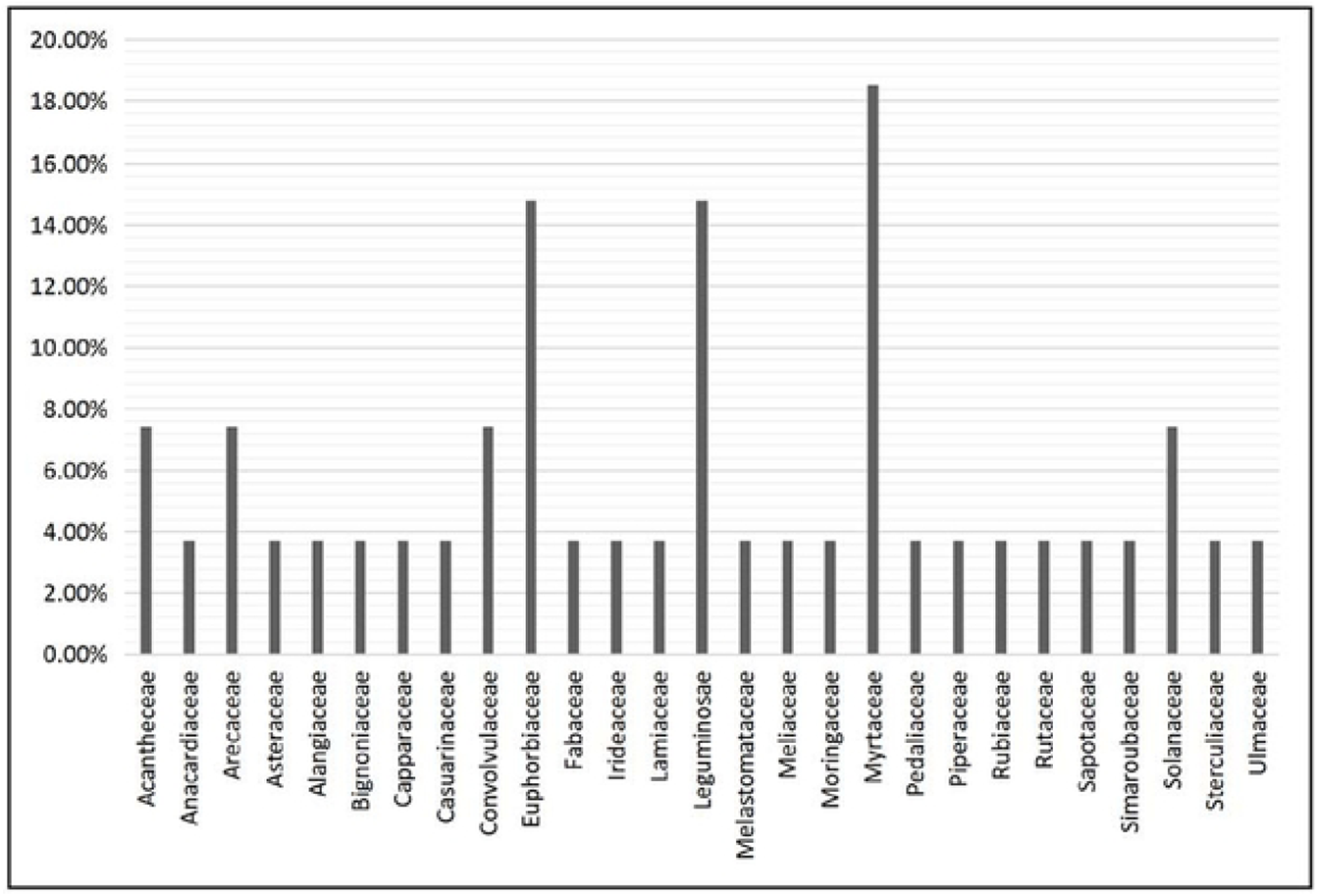
Families found in the honey samples of *Apis cerana* collected from the Eastern dry region.

**Figure 5.**
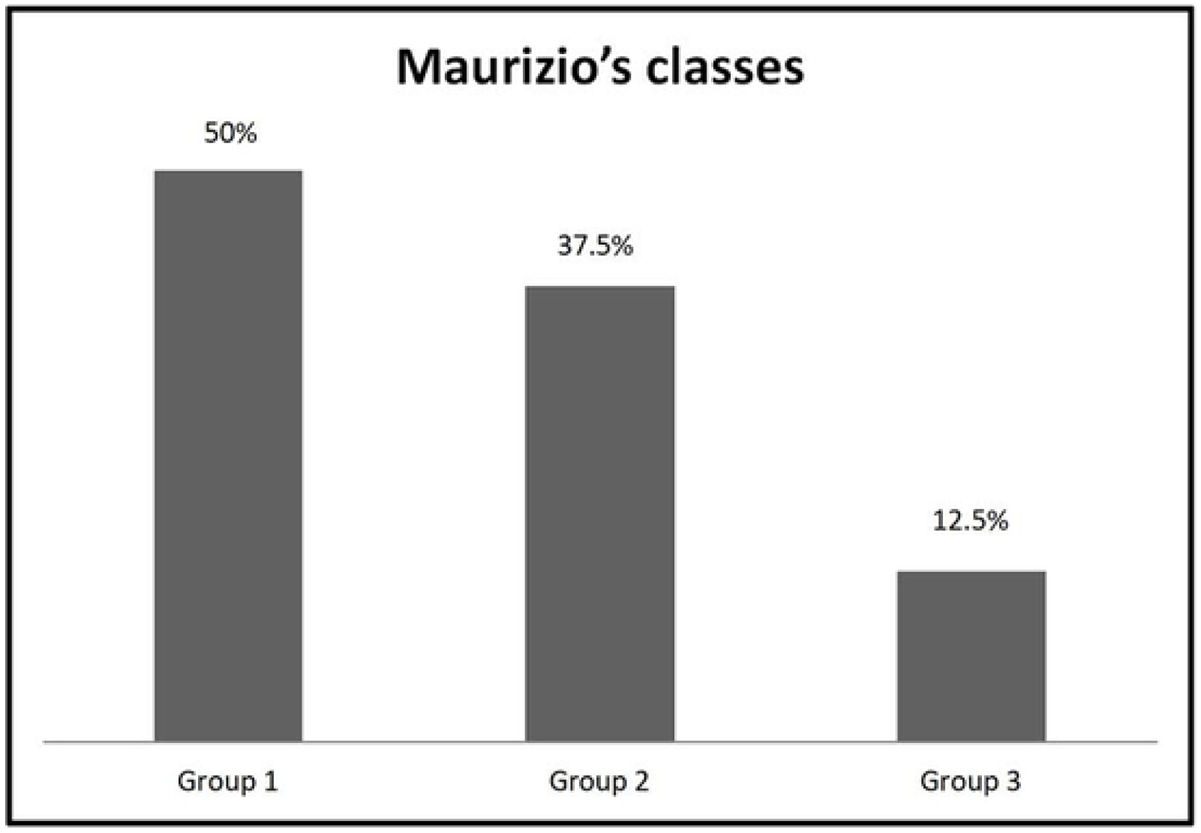
Distribution(%) of the honey samples according to Maurizio’s classes. Group I (<20,000 pollen grains per 10 g honey) found in 12 samples (50%), Group II (20,000-100,000 grains per 10 g honey) found in 9 samples (37.50%), Group Ill (100,000-500,000 grains per 10 g honey) found in 3 samples (12.5%)

Plant pollen species present in the honey samples collected across the samples were *A. auriculiformis, A. ilicifolius* L., *A. occidentale* L., *A. catechu, B. variegata* Benth., *Caesalpinia* sp., *C. viminalis* Byrnes, *C. parviflorum* Roxb., *C. zeylanica* L., *C. annuum* L., *C. equisetifolia, Citrus* sp., *C. nucifera, C. pallida* L., *Eucalyptus* sp., *E. alsinoides* L., *M. philippensis, M. zapota* (L.), *M. oleifera* L., *Ocimum* sp., *P. hysterophorus* L., *P. pinnata* L., *P. guajava* L., *S. indicum* DC., *S. glauca, S. melongena* L., *S. multiflora* Baill, *S. mahagoni* Lam., *Syzygium cumini R*.*Br*., and *Tabebuia* sp., and plant belonging to the family Sterculiaceae, Iridaceae, Myrtaceae, Alangiaceae, Acantheceae and Euphorbiaceae (Table 2, Figure 10-14).

**Figure 6:**
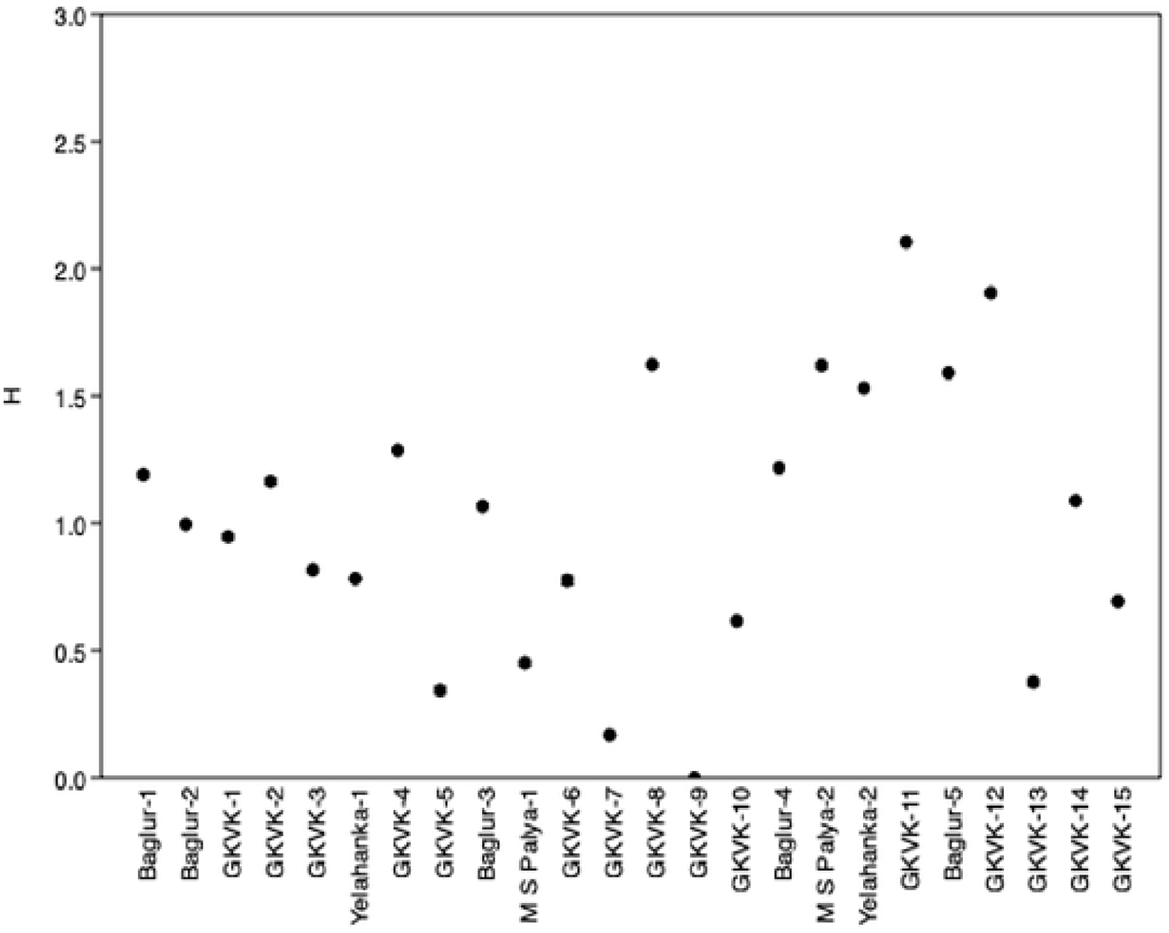
Diversity indices of the pollen samples

**Figure 7:**
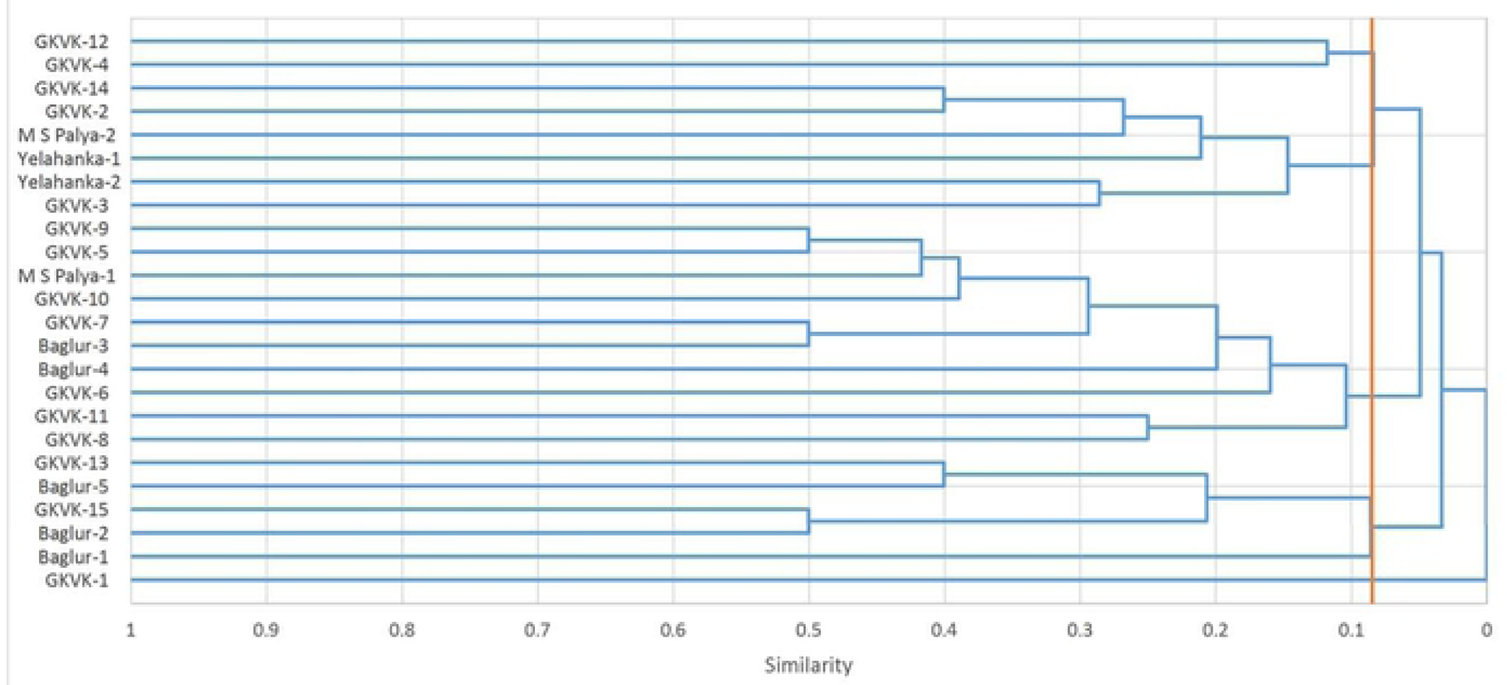
Cluster analysis of absolute pollens collected from the eastern dry zone

**Figure 8:**
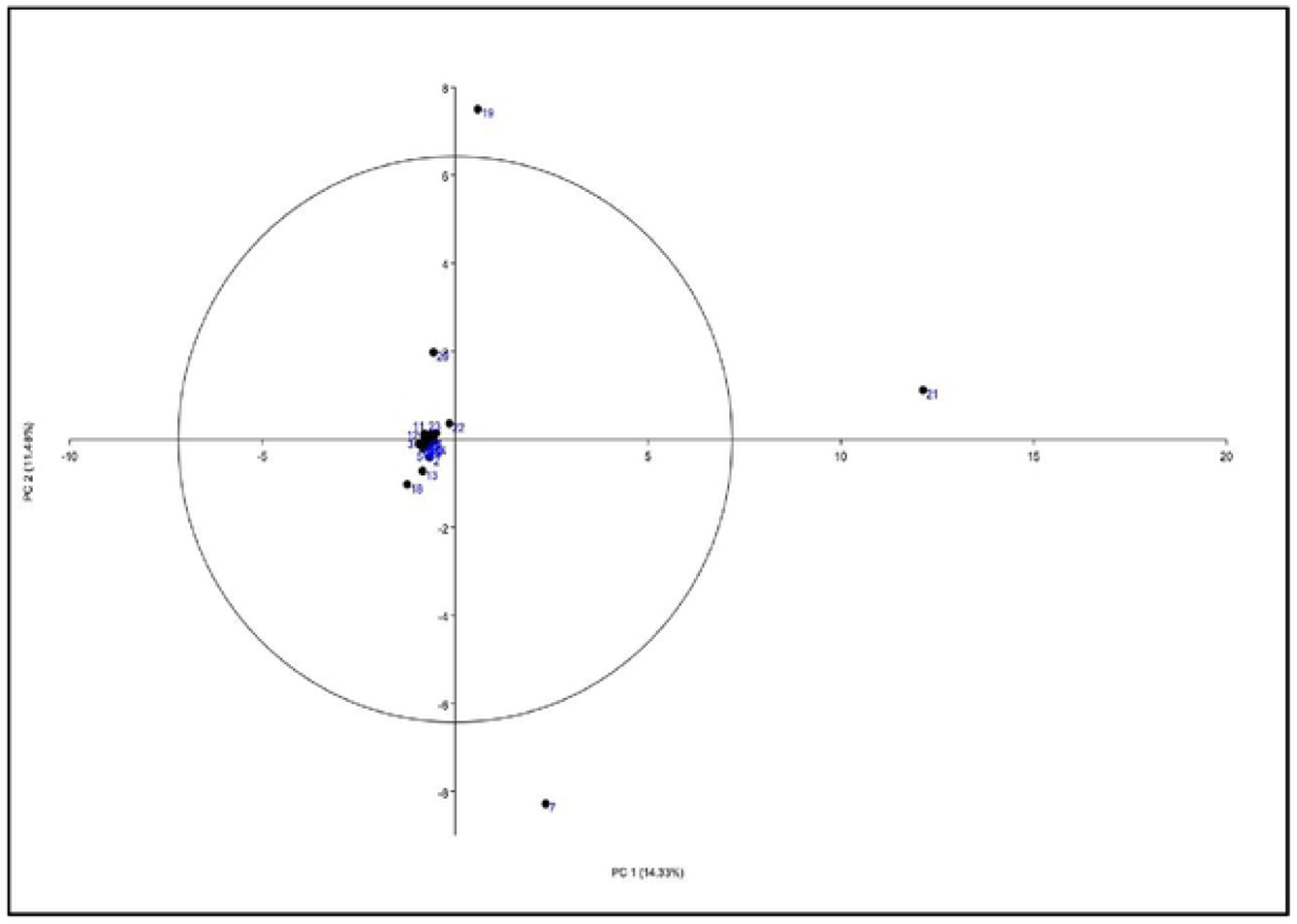
Principal component analysis using absolute pollen count

**Figure 9:**
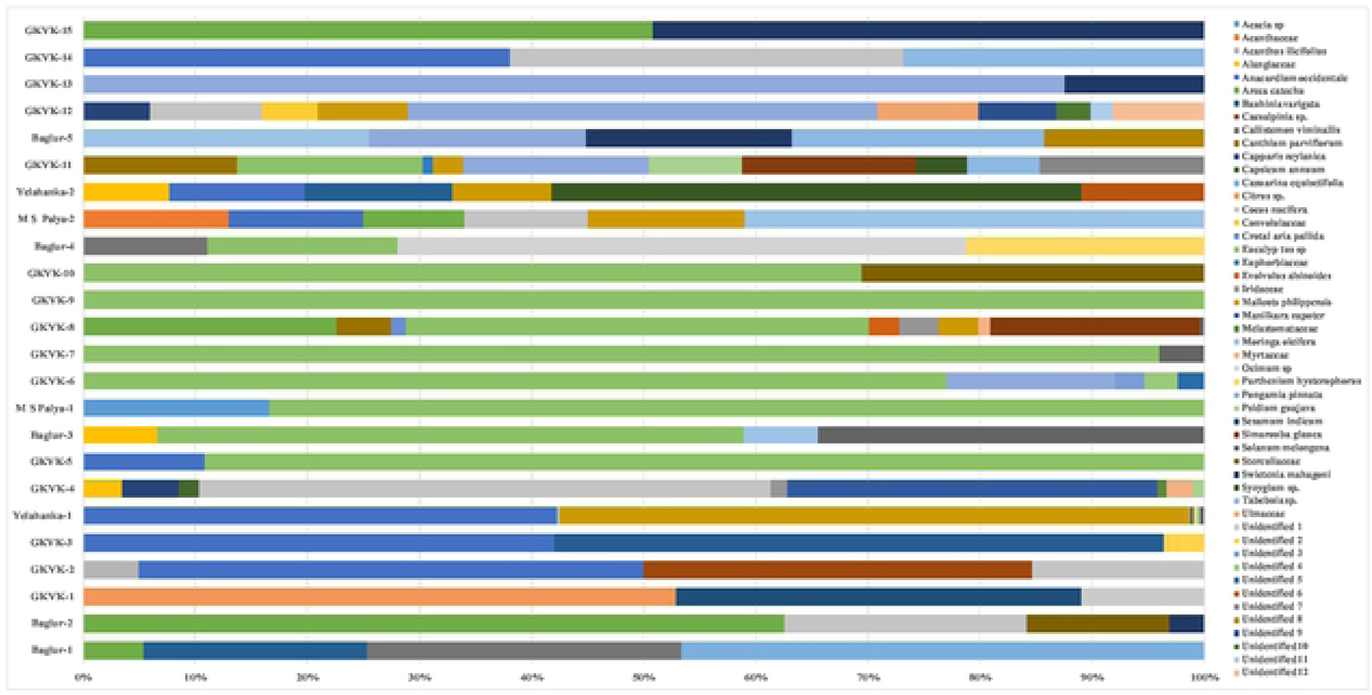
Percent frequency of pollens recorded trough melissopalynological study

**Figure. 10.**
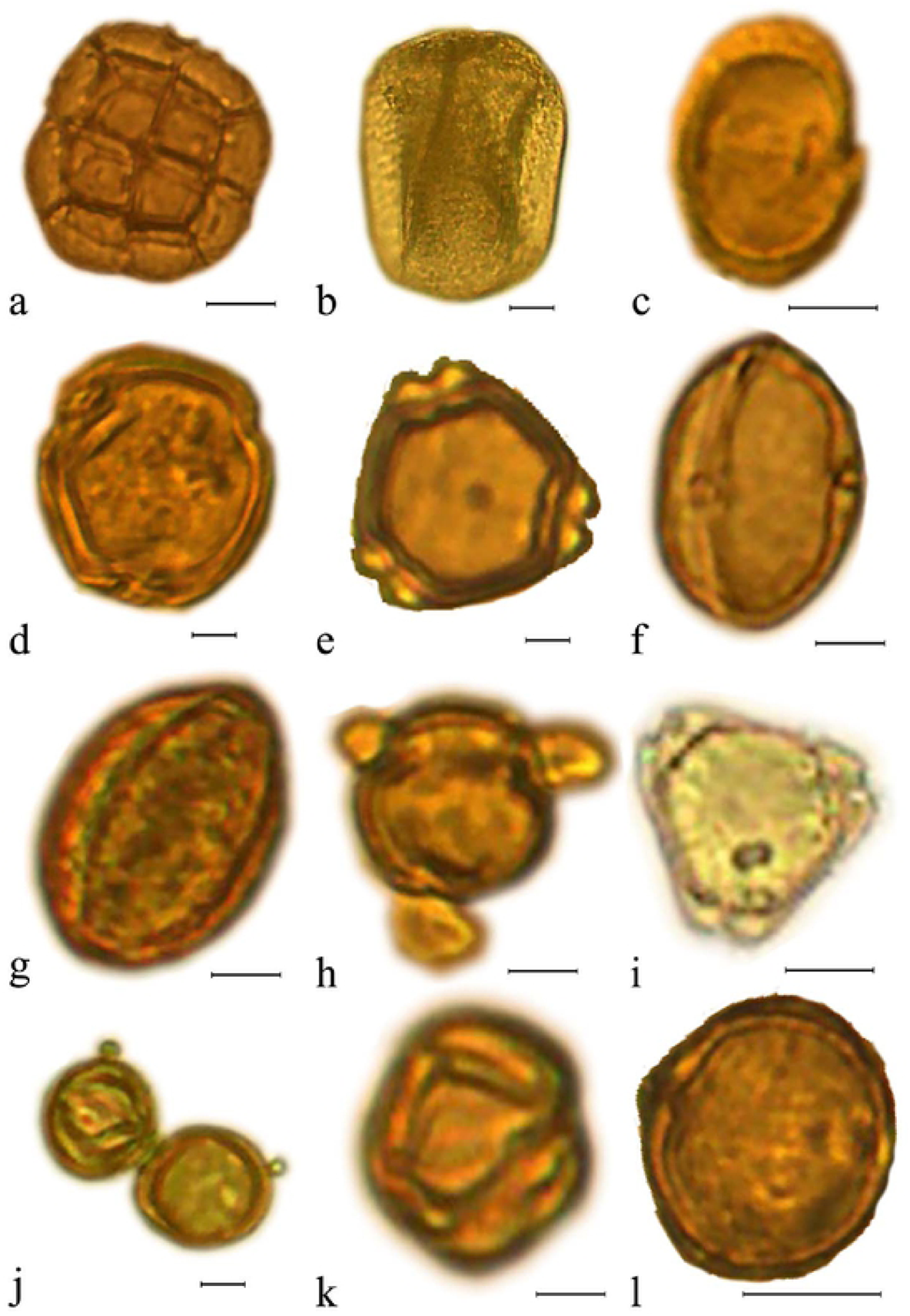
Photomicrographs of pollen grains from the honey samples. a. *Acacia* sp. L’Her. (C.Mart.), b. *Acantheceae*, c.*Acanthus ilicifolius* L, d. *Alangiaceae*, c.*Anacardium occidentale* L., f. *Areca catechu* L., *g. Bauhinia varigata* L., h. *Caesalpinia* sp. i. *Callistemon viminalis* Byrnes,j. *Canthium parviflorum* Roxb, k. *Capparis zeylanica* L., 1. *Capsicum annuum* L.

**Figure 11.**
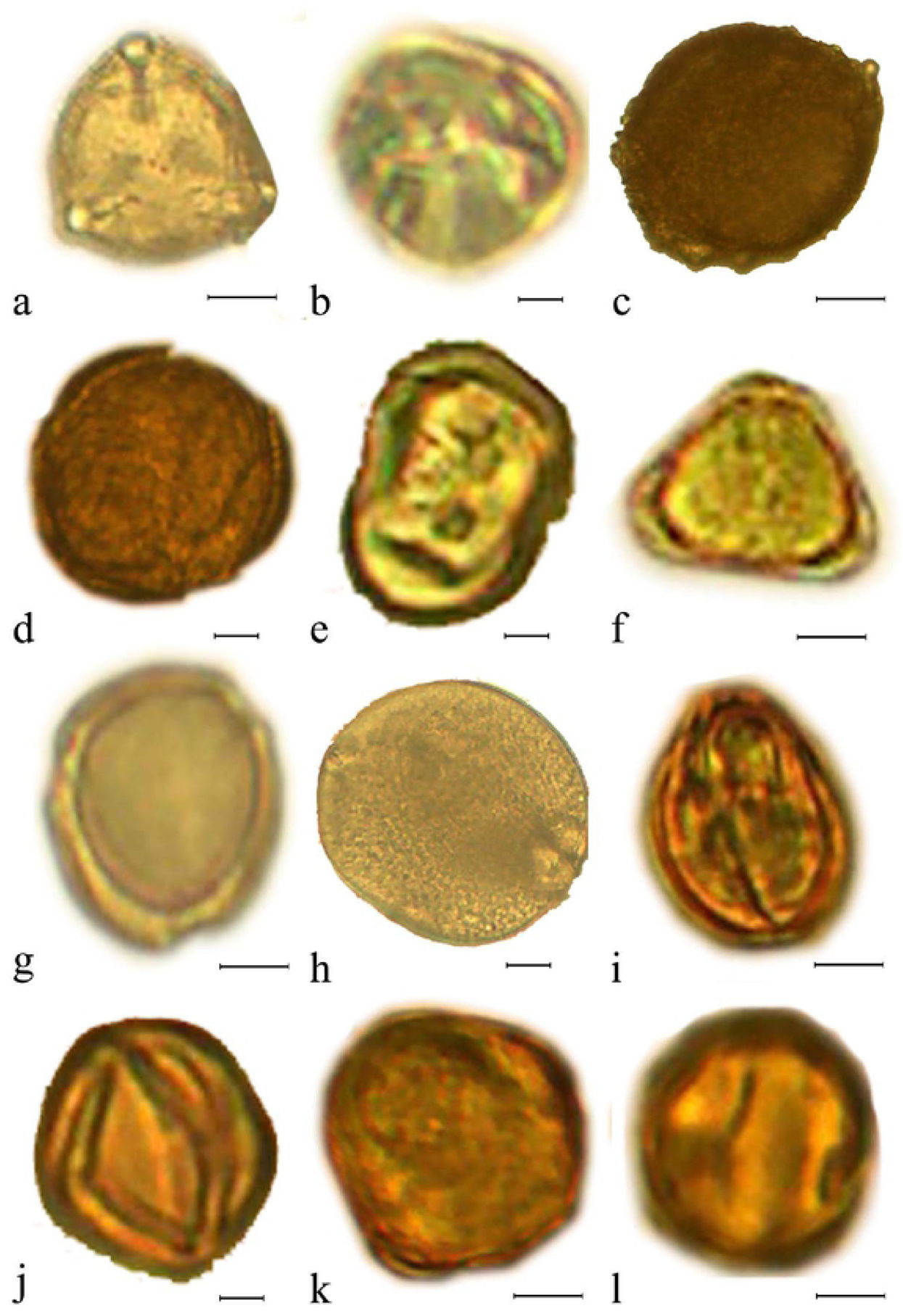
Photomicrographs of pollen grains from the honey samples (Continued). a. *Casurina equisetifolia*, b. *Citrus* sp., c. *Cocos nucifera* L., d. *Convolulaceae*, e. *Crotalaria pallida* L., *Eucalyplus* sp., *g. Euphorbiaceae, h. Evolvulus alsinoides* L., i-j. *lrideaceae*, k. *Mallotus philippensis* Lour., 1. *Manilkera zapota*.

**Figure 12.**
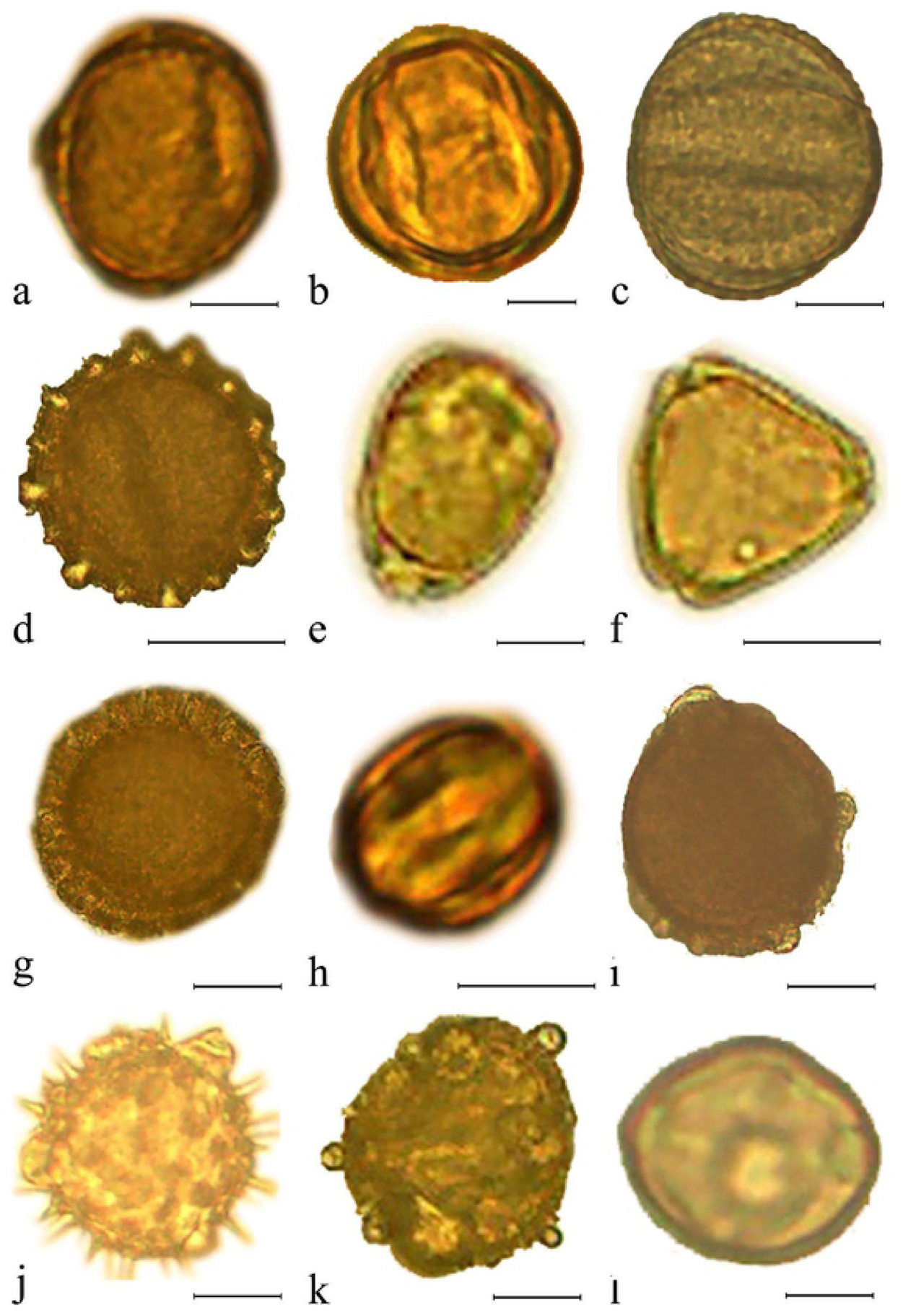
Photomicrographs of pollen grains from the honey samples (Continued). a. Melastomataccae, b. *Moringa oleifera* L., c. *Ocimum* sp. L., d. *Parthenium hysterophorus* L., c. *Pongamia pinnata* L., f. *Pisidium guajava* L., g. *Sesamum indicum* DC., h. *Simarouba glauca* DC., i. *Solanum melongena* L., J. *Sterculaceae*, h. *Suregada multiflora* Baill,j. *Swietenia mahagoni* Lam

**Figure 13.**
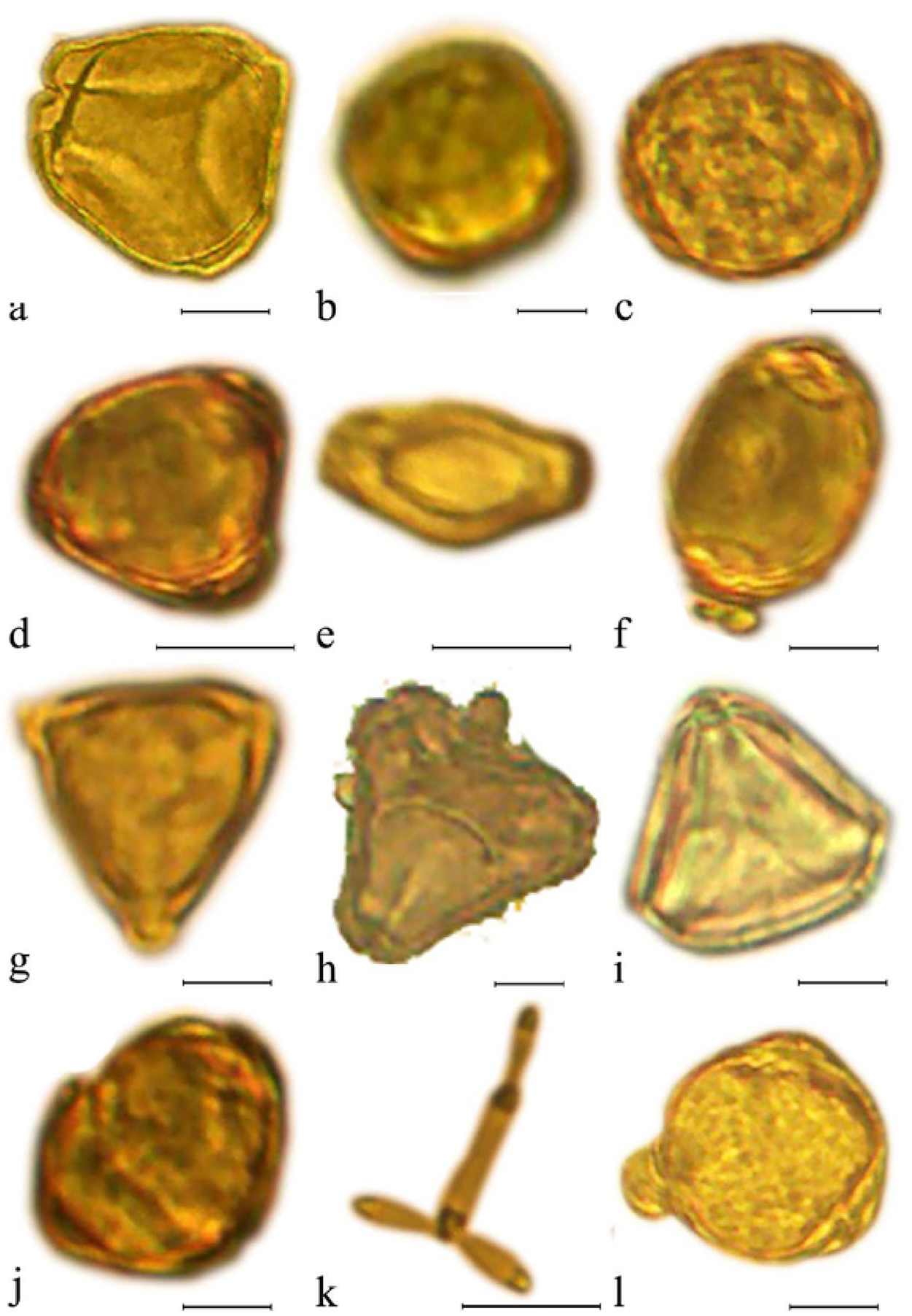
Photomicrographs of pollen grains from the honey samples (Continued). a. *Syzygium* cumini R.Br., b. *Tabebuia* sp. L., c. *Ulmaceae*, d-k. Unidentified

**Figure 14.**
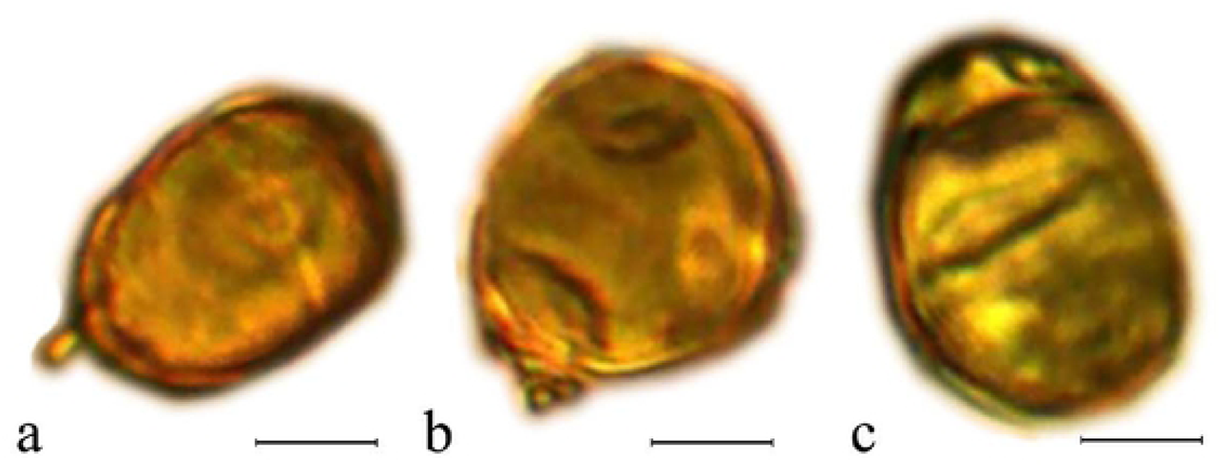
Photomicrographs of pollen grains from the honey samples (Continucd). a-c Unidentified pollen

The floral calendar of the melisopalynological studies (Figure 2) reveals that one or the other flora is available throughout the year in the Eastern Dry Zone of Karnataka. Samples collected in October, November, December, and January were rich in pollens of *Eucalyptus*. Similarly, samples collected in January, February to June had forest pollen species *viz*., *S. mahagoni, C. parviflorum, S. glauca, Eucalyptus* sp., *M. oleifera, S. cumini R*.*Br*., *Tabebuia* sp., *P. pinnata, A. occidentals, C. nucifera, A. catechu, M. Philippensis, B. variegata, P. guajava, C. zeylanica*, and plants belonging to the family *Acanthaceae, Convolvulaceae, Alangiaceae, Euphorbiaceae, Ulmaceae*.

To study the pollen diversity in the honey samples collected from different locations, Shannon diversity was determined. The diversity indices varied between 0 to 2.11 (Figure 6). Among the 24 different sources, the highest Shannon diversity was recorded in GKVK-11 followed by GKVK-12. The least diversity was recorded in GKVK-9 followed by GKVK-7. At 0.09 similarity co-efficient, data was classified into five clusters (Figure 7). Cluster 1 had an only sample (GKVK 1) which was unique since pollens (*Citrus sp*. and *Sesamum indicum*) found in this sample did not coincide with any other samples. Cluster 2 consisted of honey samples collected from locations viz., Baglur 1, Baglur 2, GKVK 15, Baglur 5, and GKVK 13. This cluster was mainly classified based on the presence of *Swietenia mahagoni*. The cluster 3 was the largest cluster comprising of 11 samples in which 81.82 % were unifloral. Cluster 4 comprised of six location and their clustering in the same group is mainly due to similarity of tree species viz., *Anacardium occidentale* L., *Bauhinia variegata* L., *Cocos nucifera* and *Mallotus philippensis*. The cluster five had two samples (GKVK 4 and GKVK 12). The Principal Component analysis revealed that most of the samples grouped into a single cluster except 7, 19, 20, and 21 which were placed away from the origin (Figure 8). The absolute pollen count and the frequency of each taxon are presented in Figure 9.

## Discussion and Conclusion

The presence of pollens in honey symbolized the bee foraging plants. Among several flowering plants bee forage specific plants (Dimou, 2007). The diversity of pollen grains in honey varied with locations and availability of bee flora (Song *et al*., 2014). The present study provides a vision on the pollen spectrum of the Eastern dry zone of Karnataka. Overall, 51 pollen taxa from 24 honey samples were recorded from the study. A total of 80 pollens were recorded from 42 samples in Tropical South India and 61 pollen from 19 samples from the central region of Shanxi (Ponnuchamy *et al*., 2014; Song *et al*., 2012). The pollens from the Eastern dry zone of Karnataka are *Callistemon viminalis* Byrnes, *Areca catechu* L., *Citrus* sp. L., *Mallotus philippensis* Lour., *Cocus nucifera* L., *Eucalyptus* sp. Labill., *Ocimum* sp. L., *Moringa oleifera* L., and *Pongamia pinnata* L., *Swietenia mahagoni* Lam., *Canthium parviflorum, Simarouba glauca* DC., *Eucalyptus* sp., *Syzygium cumini* R. Br., *Tabebuia* sp. L., *Acanthaceae, Anacardium occidentals* L., *Bauhinia variegata* L., *Psidium guajava* L., and *Capparis zeylanica* L. Similarly, in Brazil, *Eucalyptus* sp. and *Citrus* sp. were predominant (Barth, 1970: Ramlho *et al*., 1991). In India, honey from Uttar Pradesh predominated with pollen from *Antegonon* and *Moringa* (Nair and Singh, 1974), *Rumex* sp., *Nephelium* sp., and members of Myrtaceae, Liliaceae, Rosaceae, and Euphorbiaceae (Sharma and Nair, 1965; Gaur and Nanwani, 1989). Honey from Himachal Pradesh had a preponderance of *Brassica, Adathoda, Clematis, Mussenda*, and *Helianthus* sp. (Singh *et al*., 1994). Honey from Andhra Pradesh revealed that *Sapindus, Eucalyptus, Anacardium*, and *Cleome* were major pollen types (Kalpana and Ramanujan, 1997). In Karnataka, honey samples were having pollens from *Cocos, Eucalyptus, Schefflera*, and *Mimosa* (Singh and Suryanarayana, 1990). Also, the Samples collected in October, November, December, and January were rich in pollens of *Eucalyptus*. Many *Eucalyptus* species offer pollen and nectar to the pollinators (House, 1997). In the many tropical to subtropical regions, where *Eucalyptus* species have been planted, and often become naturalized, they have become important, or even dominant, nectar sources for beekeeping in southern Asia (Chauhan *et al*., 2017), in South America (Daners, 1998; Bonilla *et al*., 2016), in Africa (Carroll, 2006) and other tropical regions (Rasoloarijao, 2014), and many countries within the Mediterranean Basin (Seijo *et al*., 2003; Terrab *et al*., 2003; Feás *et al*., 2010). Similarly, in January, February and March had forest pollen species *viz*., *Pongamia, Syzygium, Santalum, Neem, Mahagoni*, etc. The presence of pollen in the honey of particular plant species during different months is related to the blooming of that particular plant species from which the bees collected the pollen during foraging activity (Joshi *et al*., 1998). They have reported that the major honey sources were *Bombax, Lannea, Limonia, Moringa, Peltoforum, Pongamia, Syzygium*, and *Tamarindus* during major honey flow season (February-July) and the pollens of *Eucalyptus* and *Alternanthera* were predominant during minor honey flow season (September-December).

The results of this work also reveal that 62.5% of the honey samples are unifloral and the remaining 37.5% are multifloral and the floral type of honey was confirmed by the absolute pollen count. From the Leon and Palenica provinces out of 89 honey samples 51.69% was unifloral (Herrero *et al*., 2001). The quality and quantity of the bee pollen produced and the floral type help in increasing the market value of honey.

A floral calendar is developed with month-wise bee foraging plants with the classifying groups viz., major pollens, secondary pollen, and minor pollen types. This will help farmers and beekeepers to educate about the flora available and establish the apiary.

Shannon diversity indices are used to classify the samples based on their spatial location and to gather knowledge about the dynamics of preference toward the foraging plant (Ponnuchamy *et al*., 2014). High values of Shannon diversity indices in samples GKVK-11 and GKVK-12 indicates rich nectar and pollen sources in GKVK location in March. The Shannon-Weaver diversity index was varying from 1.79 to 2.21 in the samples from the Central region of Shanxi, North China (Song *et al*., 2012). Some of the locations had similar pollen compositions based on which the cluster analysis is plot. From the statistical analysis of cluster analysis, the samples are grouped into five clusters. This result is in accordance with the four clusters formed out of 89 samples which were collected from the Leon and Palenica provinces (Herrero *et al*., 2001). This is based on the common pollen taxa recorded in different months. *Eucalyptus* sp. L was predominant in samples of cluster 4. GKVK 1 recorded unique compared to other cluster; this may be due to *Sesamum* and *Citrus*, generally, *Sesamum* isn’t grown in the Eastern dry zone of Karnataka and few experimental plots of GKVK campus of UAS, Bangalore, had *Sesamum* which reflected in GKVK 1 sample. Majority samples in cluster 3 are unifloral and cluster 4 are multifloral. Based on this study, honey samples are grouped according to their botanical origin.

The grouping in PCA might be due to pollen diversity, the season of collection and abundance or dearth period (Sekhar, 2000). This was reflected in PCA plots wherein a few samples (7, 19, 20, 21) were distributed away from the origin (Fig. 8). Since most of the tree species in the study location bears flowers in February and March, the honey bees had the opportunity to explore the diverse flora and collect pollens of different tree species. It was also observed that samples that had pollens from more than 5 species might have been placed away from the origin. It can be observed that the time of sampling would have affected the pollen variability in the honey sample (Marc, 2012; Raja, 2012). Among these samples, 75% were multifloral and were sampled during March.

Overall, this study provides useful data regarding the favoured plant and the foraging preference of *Apis cerana* from the Eastern dry zone of Karnataka, which will help beekeepers to develop the apiary. It provides information regarding the agricultural crop which gets pollinated and can be utilized for pollination purposes. This study will in-turn contribute to the conservation of bees, apiary development and leads to sustainable honey production. Also, this will improve the socioeconomic status of the farmers and beekeepers.

## Acknowledgement

We are sincerely thankful to Dr. Bhat, Mrs. Eshwarappa G. C., Dr. Vijayakumar KT for providing valuable guidance throughout the project.

